# Axillary Microbiota Compositions from Men and Women in a Tertiary Institution-South East Nigeria: Effects of Deodorants/Antiperspirants on Bacterial Communities

**DOI:** 10.1101/2020.03.11.986364

**Authors:** Kingsley C Anukam, Victoria Nmewurum, Nneka R Agbakoba

## Abstract

The axillary skin microbiota compositions of African populations that live in warm climate is not well studied with modern next-generation sequencing methods. To assess the microbiota compositions of the axillary region of healthy male and female students, we used 16S rRNA metagenomics method and clustered the microbial communities between those students that reported regular use of deodorants/antiperspirants and those that do not. Axillary skin swab was self-collected by 38 male and 35 females following uBiome sample collection instructions. Amplification of the V4 region of the 16S rRNA genes was performed and sequencing done in a pair-end set-up on the Illumina NextSeq 500 platform rendering 2 × 150 base pair. Microbial taxonomy to species level was generated using the Illumina Greengenes database. 26 phyla were identified in males with *Actinobacteria* as the most abundant (60%), followed by *Firmicutes* (31.53%), *Proteobacteria* (5.03%), *Bacteroidetes* (2.86%) and others. Similarly, 25 phyla were identified in females and *Actinobacteria* was the most abundant (59.28%), followed by *Firmicutes* (34.28%), *Proteobacteria* (5.91%), *Bacteroidetes* (0.45%) and others. A total of 747 genera were identified, out of which 556 (74.4%) were common to both males and females and 163 (21.8%) were exclusive to males while 28 (3.8%) were exclusive to females. *Corynebacterium* (53.89% vs 50.17%) was the most relative abundant genera in both male and female subjects, followed by *Staphylococcus* (19.66% vs 20.90%), *Anaerococcus* (4.91% vs 7.51%), *Propionibacterium* (1.21% vs 1.84%). There was a significant difference (*P*=0.0075) between those males that reported regular use of antiperspirant/deodorants and those that reported non-use of antiperspirants/deodorants in the relative abundance of *Corynebacterium* (68.06% vs 42.40%). Higher proportion of *Corynebacterium* was observed in male subjects than females, while more relative abundance of *Staphylococcus* was found in females than males. This study detected *Lactobacilli* in the axilla of over 82% of female and over 81% of male subjects, though in low relative abundance which suggests that *Lactobacillus* taxa might be considered as part of the normal axillary bacterial community. The study also revealed that the relative abundance of *Corynebacterium* (68.06% vs 42.40%) was higher in those that reported regular use of deodorants/antiperspirants.

## INTRODUCTION

The bacterial microbiota compositions of the axillary skin of African people is less well studied with the modern next-generation sequencing technology resulting in little or poor knowledge on the microbial communities that could be mined for diagnostic and therapeutic purposes. Previous studies on the axillary microbiota have relied on culture-dependent methods whereby *Staphylococci* or *Corynebacteria* genera have consistently been incriminated (Jackman, 1983; Taylor, et al., 2003). This is due to the fact that culture methods utilize artificial media that can support the growth of these bacteria leading to underestimation of other microbes present on the body site (Kong and Segre, 2012). One culture-dependent study that was conducted in people affected by Albinism and those with normal pigmented skin in Northern Tanzania showed that *Staphylococcus* was the commonest microorganism isolated in over 90% of the samples (Kiprono et al., 2012). In the last decade, the use of next-generation sequencing approach has revealed an avalanche of microbial communities that inhabit the axillary region showing the predominance of *Staphylococci, Corynebacteria, Anaerococcu*s and *Peptoniphilus* (Egert et al., 2011; Callewaert et al., 2013; Troccaz et al., 2015).

Body malodour is the most common reason human adults generally use deodorants or antiperspirants in order to obtain an appealing body odour or to mask and reduce sweat from the apocrine glands. Bacteria present in the skin are responsible for body odour, whereby sweat components which are odourless are broken down to odour-causing substances such as steroid derivatives, short volatile branched-chain fatty acids and sulphanylalkanols. In the underarm or axilla, malodour arises due to biotransformation by the microbiota of dipeptide-conjugated thioalcohols, particularly S-[1-(2-hydroxyethyl)-1-methylbutyl]-(l)-cysteinylglycine (Cys-Gly-3M3SH) (Bawdon et al., 2015). Most students in tertiary institutions around the world are conscious of body odour and application of deodorants have recorded corresponding influence on the species diversities of the axillary microbiome (Callewaert et al 2013). In Western societies, over 95% of the young adult population are concerned about their personal hygiene and are less tolerant toward unpleasant body odour and they make use of underarm deodorants and antiperspirants(Callewaert et al., 2014). In the same way, the adult population in Africa and particularly students, utilize deodorants with the perception of increased social confidence and improvement in the quality of life (Pierard et al., 2003). We do not have documented information on the social predictors that motivate university students in Nigeria on the regular use of antiperspirants/deodorants but it is believed to be a common phenomenon as marketers advertise such products with hype, brazenly. In this study we hypothesized that the relative abundance of bacterial communities in adult male students may be different from adult female students. The objectives of this study are two folds: first to determine the microbiota compositions of the axillary region of healthy male and female students using 16S rRNA metagenomics method and second to separate the microbial communities between those students that reported regular use of deodorants/antiperspirants and those that do not.

## MATERIALS AND METHODS

### Ethics Review Committee Approval

This study was carried out in accordance with the recommendations of the ethic review committee of the Faculty of Health Sciences, Nnamdi Azikiwe University. All subjects gave written informed consent in accordance with the Declaration of Helsinki.

### Study Participants and Sample Collections

A total of 100 participants comprising of 50 male and 50 female students from the Faculty of Health Sciences & Technology, Nnamdi Azikiwe University, Nnewi Campus were recruited in the study. The selection criteria involved those with no history of dermatological disorders or other chronic medical disorders and with no current skin infections. Participants were between the ages of 17 years old to 35 years old. They provided signed informed consents. Socio-demographic data, skin health or disease history and regular use of deodorant/antiperspirants were obtained from the participants through the administered questionnaires. Skin (Axilla) sample was self-collected following uBiome® sample collection instructions. A moistened sterile cotton swab (uBiome) was thoroughly swabbed for 20 seconds in the axillary region to detach and absorb the microorganisms, and it was vigorously agitated for 20 seconds in a sterilized reaction vial or tube containing a lysis and stabilization buffer that preserves the DNA for transport at ambient temperatures. The tubes were sent to uBiome Inc. in California, United States America for DNA extraction and sequencing. Sequencing results were analyzed with bioinformatic tools at Uzobiogene Genomics, London, Ontario, Canada.

### DNA Extraction and Sequencing of the 16S rRNA V4 region

Bacterial DNA was extracted from the axilla swabs using an in-house protocol developed by uBiome Inc. Briefly, samples were lysed using bead-beating, and DNA was extracted in a class 1000 clean room by a guanidine thiocyanate silica column-based purification method using a liquid-handling robot. PCR amplification of the 16S rRNA genes was performed with primers containing universal primers amplifying the V4 region (515F: GTGCCAGCMGCCGCGGTAA and 806R: GGACTACHVGGGTWTCTAAT) as previously described (Caporaso et al, 2011). In addition, the primers contained Illumina tags and barcodes. DNA samples were barcoded with a unique combination of forward and reverse indexes allowing for simultaneous processing of multiple samples. PCR products were pooled, column-purified, and size-selected through microfluidic DNA fractionation. Consolidated libraries were quantified by quantitative real-time PCR using the Kapa Bio-Rad iCycler qPCR kit on a BioRad MyiQ before loading into the sequencer. Sequencing was performed in a pair-end modality on the Illumina NextSeq 500 platform rendering 2 × 150 bp pair-end sequences. The sequencer has a flow cell with four lanes. This means that each sample was read in four different lanes (L001 to L004), and each produced forward (R1) and reverse (R2) reads.

### Metagenomics Sequence Analysis

Raw sequence reads were demultiplexed using Illumina’s BCL2FASTQ algorithm. Reads were filtered using an average Q-score > 30. The 8 paired-end sequence FASTQ reads for each sample were imported into MG-RAST pipeline for quality check (QC). Artificial replicate sequences produced by sequencing artifacts were removed following Gomez-Alvarez, et al., (2009) protocol. Any human host specific species sequences were removed using DNA level matching with bowtie (Langmead et al. 2009) and low-quality sequences were removed using a modified DynamicTrim method by Cox et al. (2011). Quantitative Insights into Microbial Ecology (QIIME) pipeline was used for 16S rRNA recognition. Sequences were pre-screened using QIIMEUCLUST algorithms for at least 97% identity to ribosomal sequences from the RNA databases. Reads passing all above filters were aligned to the database of 16S rRNA gene sequences. Microbial taxonomy to species level was generated using the Illumina BaseSpace Greengenes database.

## RESULTS

We hereby present the 16S rRNA dataset of the axillary skin microbiome compositions from the students. Out of 100 axillary swab samples that were collected from male and female students, 38 male and 35 female samples passed quality check and were analyzed with bioinformatics tools. On average the base pair count contains 29,129,902bp of DNA sequence and the sequence count contains 194,736 sequences ranging from 32bp to 151bp and averaging 149bp in length (std.deviation from average length 8.777). All of the sequences have unique identifications. The average GC-content is 55.770% (std.deviation 2.740) and GC-ratio 0.802 (std.deviation 0.095). Distribution of the taxonomic categories shows that the axilla of the male subjects had phyla that ranged from 5-22, Class (10-38), Order (14-80), Family (30-163), Genus (42-326), and Species (35-565) as shown in **Figure 1**.

**Figure 1:**
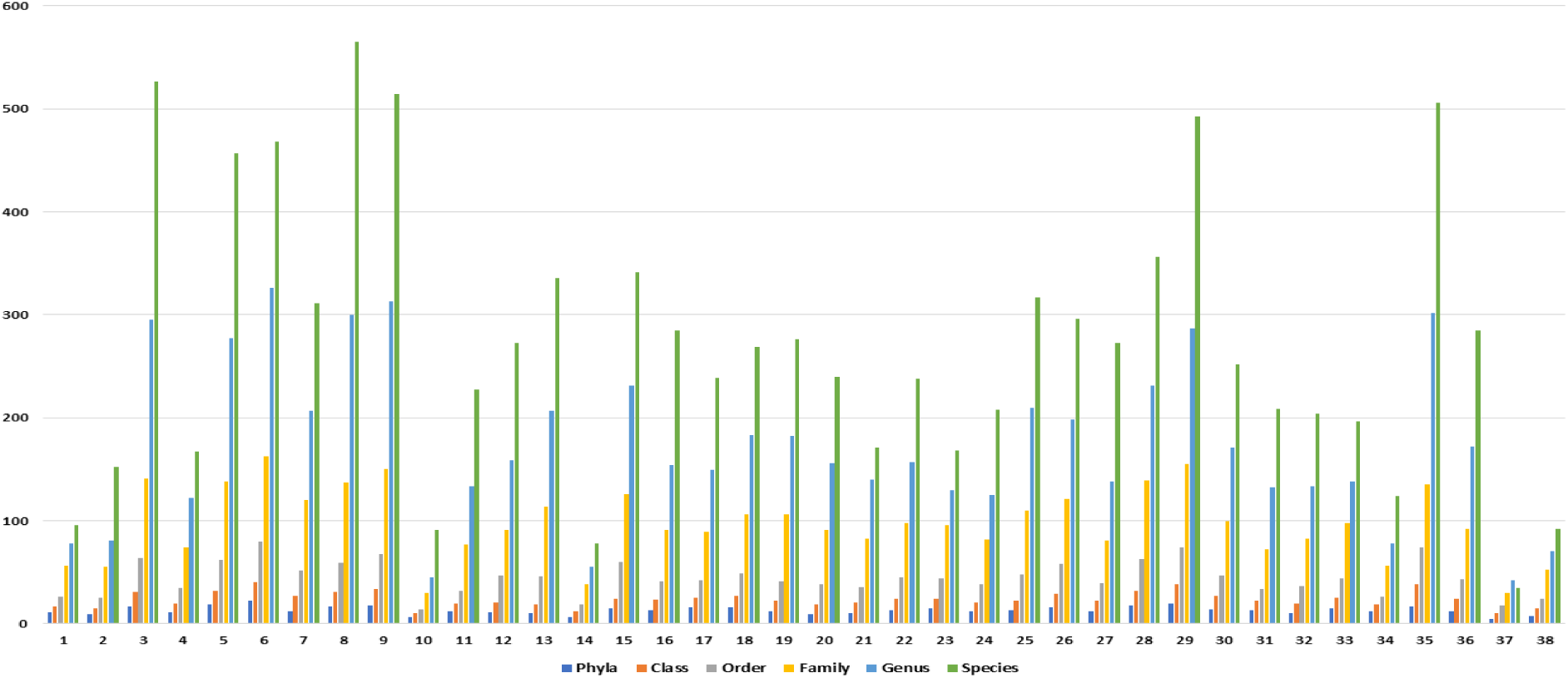
Taxonomic distribution categories in male subjects.

In contrast, the female subjects had phyla (6-18), Class (11-32), Order (17-69), Family (31-148), Genus (47-292), and Species (79-566) presented in **Figure 2**.

**Figure 2:**
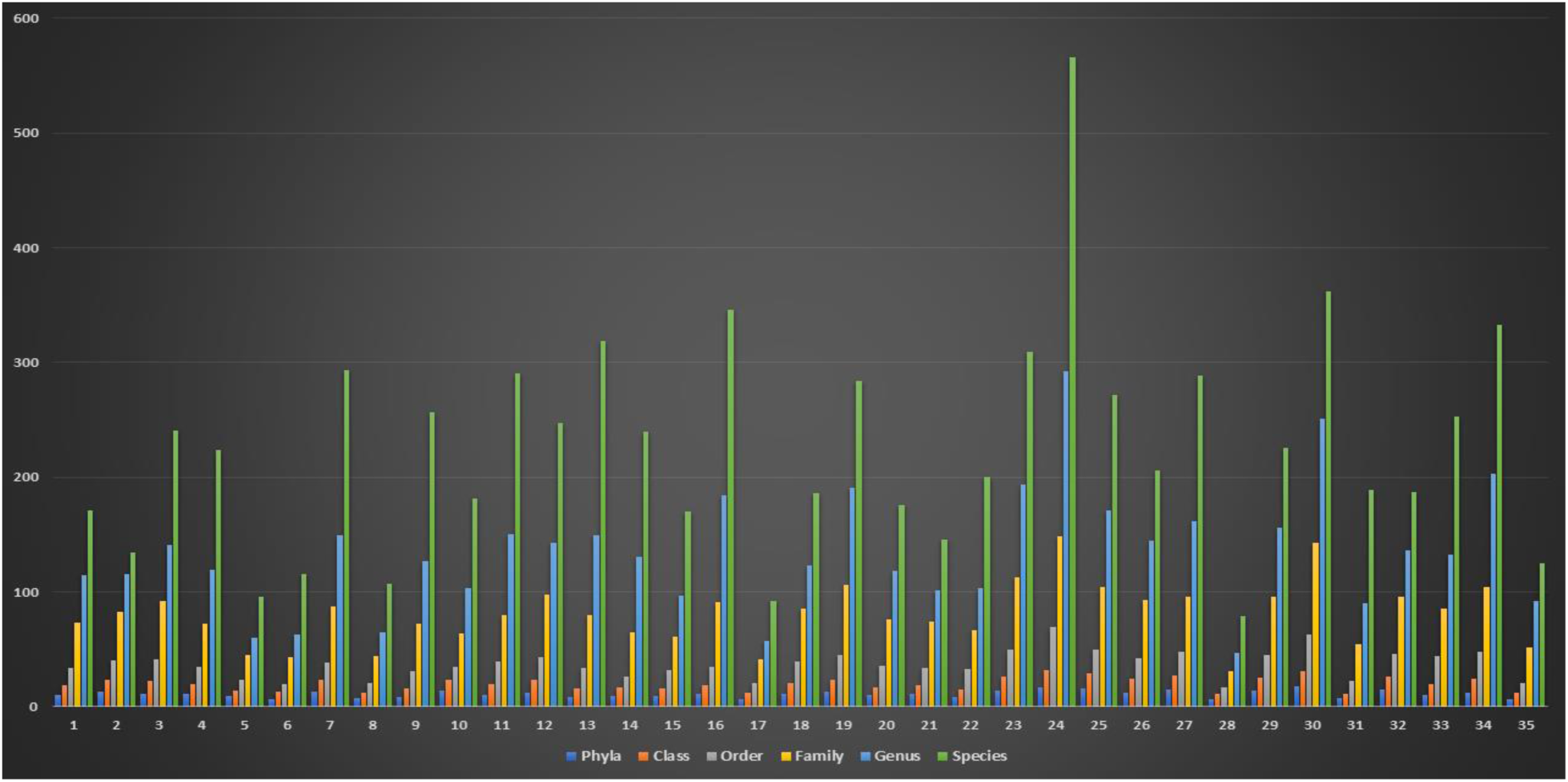
Taxonomic distribution categories in female subjects.

26 phyla were identified in males with *Actinobacteria* as the most abundant (60%), followed by *Firmicutes* (31.53%), *Proteobacteria* (5.03%), *Bacteroidetes* (2.86%) and others. Two phyla, *Fibrobacteres* and *Nitrospira*e appeared exclusive to the males. Similarly, 25 phyla were identified in females and *Actinobacteria* was the most abundant (59.28%), followed by *Firmicutes* (34.28%), *Proteobacteria* (5.91%), *Bacteroidetes* (0.45%) and others as shown in **Figure 3**. *Caldithrix*, occurred exclusively in the female samples.

**Figure 3:**
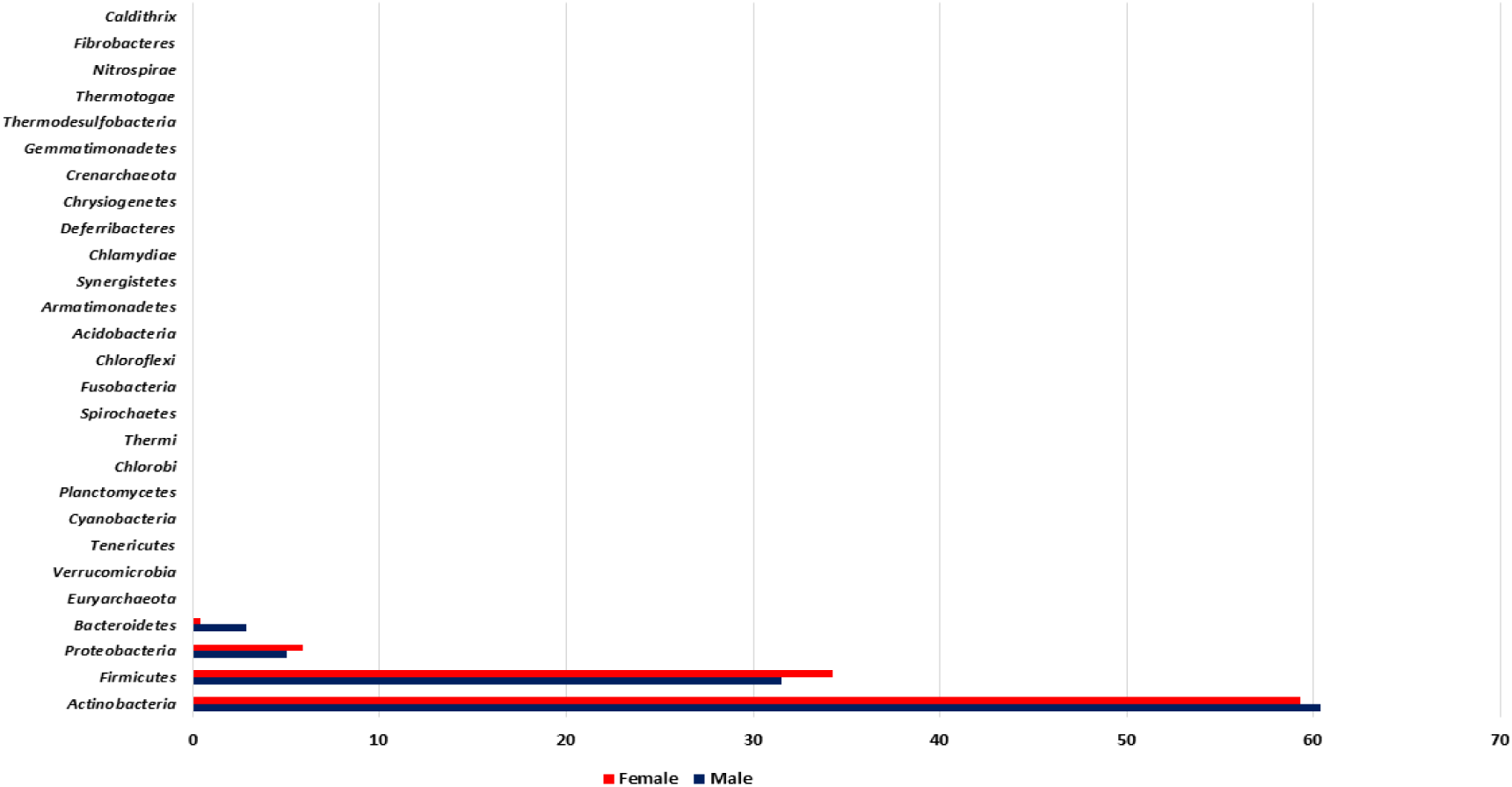
Phyla relative abundance (%) in both male and female subjects.

At the Family taxonomic level, 257 families were identified, out of which 211 were common to both male and females and 40 were exclusive to males, while 6 families were exclusive to females. Three common families including Corynebacteriaceae (56%/52%), Staphylococcaceae (20%/22%) and Clostridiaceae (6.8%/10.6%) appeared as the most abundant families in both males and females respectively. Among the exclusive families identified in females were Caldithrixaceae, Sporichthyaceae, Halothiobacillaceae, Cohaesibacteraceae, Nannocystaceae and Sulfolobaceae.

At the genera taxonomic level, a total of 747 genera were identified, out of which 556 (74.4%) were common to both males and females and 163 (21.8%) were exclusive to males while 28 (3.8%) were exclusive to females. *Corynebacterium* (53.89%) was the most relative abundant genera in males, followed by Staphylococcus (19.66%), Anaerococcus (4.91%), Propionibacterium (1.21%), Bacteroides (1.14%), Kaistella (0.85%), Faecalibacterium (0.79%), Blautia (0.76%), Acinetobacter (0.69%) and others. Similarly, *Corynebacterium* (50.17%) was the most relative abundant genera in females, followed by Staphylococcus (20.90%), Anaerococcus (7.51%), Acinetobacter (2.79%), Propionibacterium (1.84%), Enhydrobacter (1.68%), Micrococcus (1.64%), Finegoldia (1.47%), Peptoniphilus (1.08%), Exiguobacterium (1.03%), Mycobacterium (0.41%), Pseudoclavibacter (0.28%) and others as shown in **Figure 4**.

**Figure 4:**
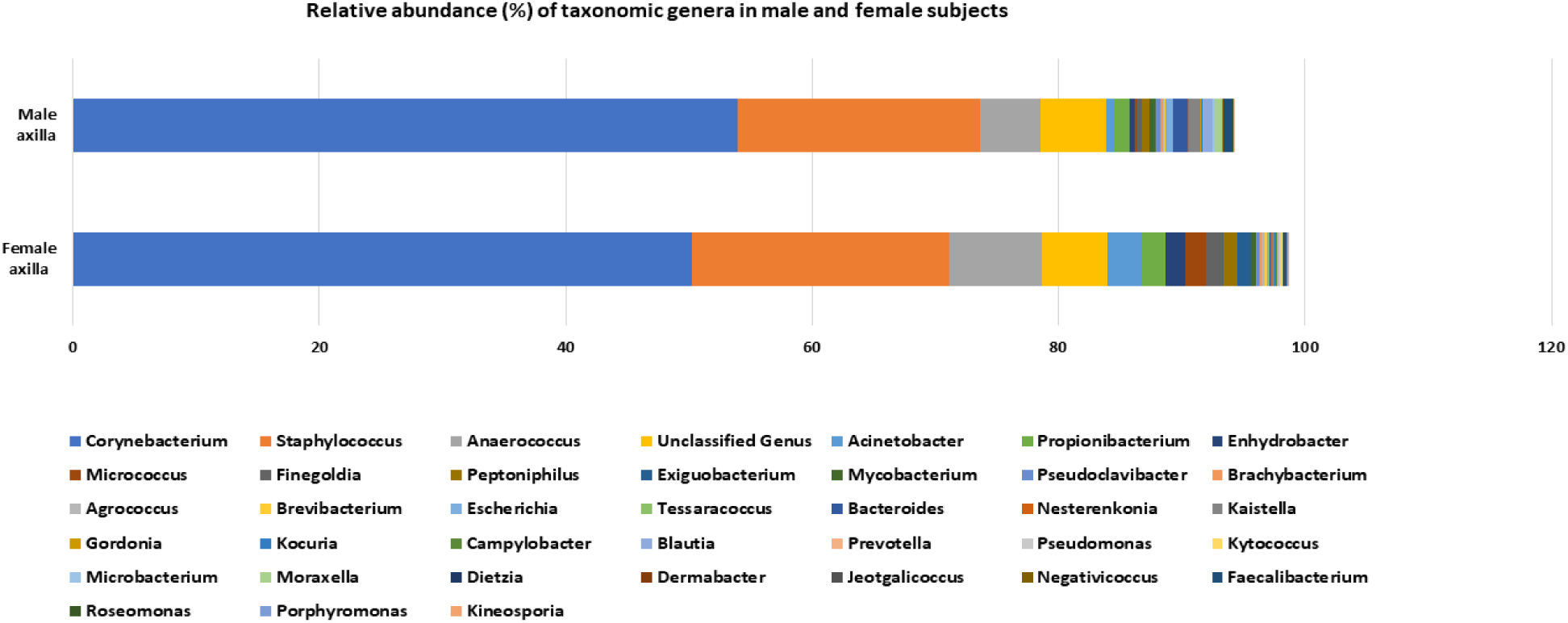
Comparative relative abundance (%) of taxonomic genera in male and female subjects.

Comparatively, there was a significant difference between male and female on the relative abundance of *Corynebacterium* (*P*=0.016), *Acinetobacter* (*P*=0.050), *Enhydrobacter* (*P*= 0.0001), *Finegoldia* (*P*=0.000013), *Micrococcus* (*P*=0.0005), and *Kaistella* (P=0.0145). The proportion of *Lactobacillus* genera found in 29/35 females was higher (0.02%) compared to 0.01% found in 31/38 of males.

At the species taxonomic level, a total of 1994 species were identified of which 1134 species were common to both male and female subjects, while 612 species were exclusively found in males (***Supplementary Table 1***) and 248 species were identified exclusively in females (***Supplementary Table 2***). Among the male subjects, *Corynebacterium appendicis* (19.86%) was the most abundant species, followed by *Corynebacterium glaucum* (6.35%), *Corynebacterium sundsvallense* (6.18%), *Corynebacterium coyleae* (5.65%), *Corynebacterium tuberculostearicum* (5.18%), *Corynebacterium tuscaniense* (4.41%), *Corynebacterium riegelii* (4.05%), *Corynebacterium imitans* (4.02%), *Anaerococcus octavius* (3.82%), *Staphylococcus haemolyticus* (2.82%) and others represented in **Figure 5**.

**Figure 5:**
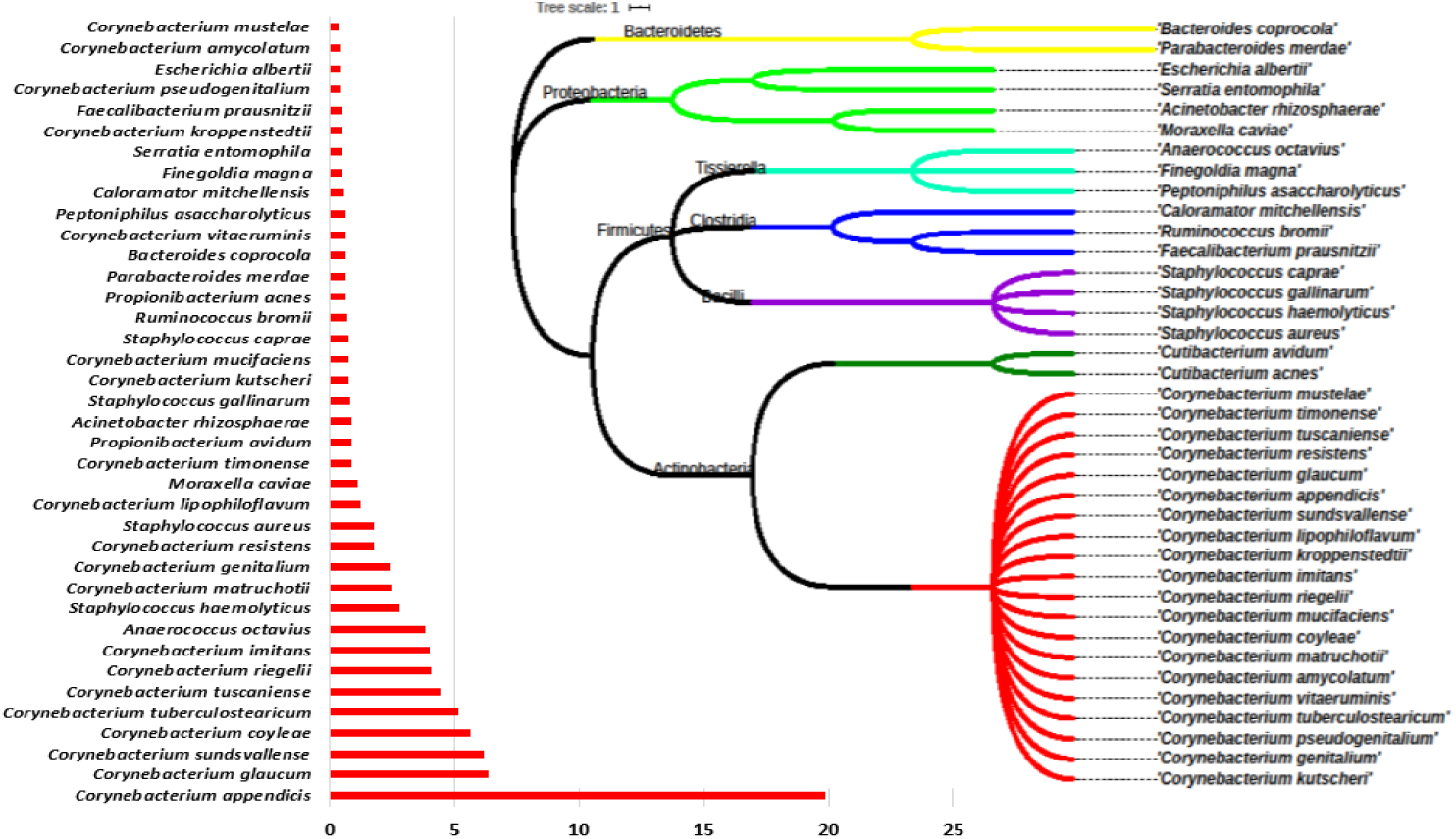
The most relative abundant (%) species in the axilla of male subjects.

In contrast, among the female subjects, *Corynebacterium tuberculostearicum* (20.80%) was the most abundant species identified in all females subjects followed by Corynebacterium coyleae (9.75%), *Corynebacterium appendicis* (5.41%), *Corynebacterium glaucum* (5.04%), *Anaerococcus octavius* (4.39%), *Corynebacterium mucifaciens* (3.38%), *Staphylococcus haemolyticus* (3.04%), *Corynebacterium kroppenstedtii* (2.86%), *Finegoldia magna* (2.80%), *Micrococcus yunnanensis* (2.55%), *Staphylococcus aureus* (2.51%) and others as shown in **Figure 6**. Comparative relative abundance of *Corynebacterium* species is presented in **Figure 7**.

**Figure 6:**
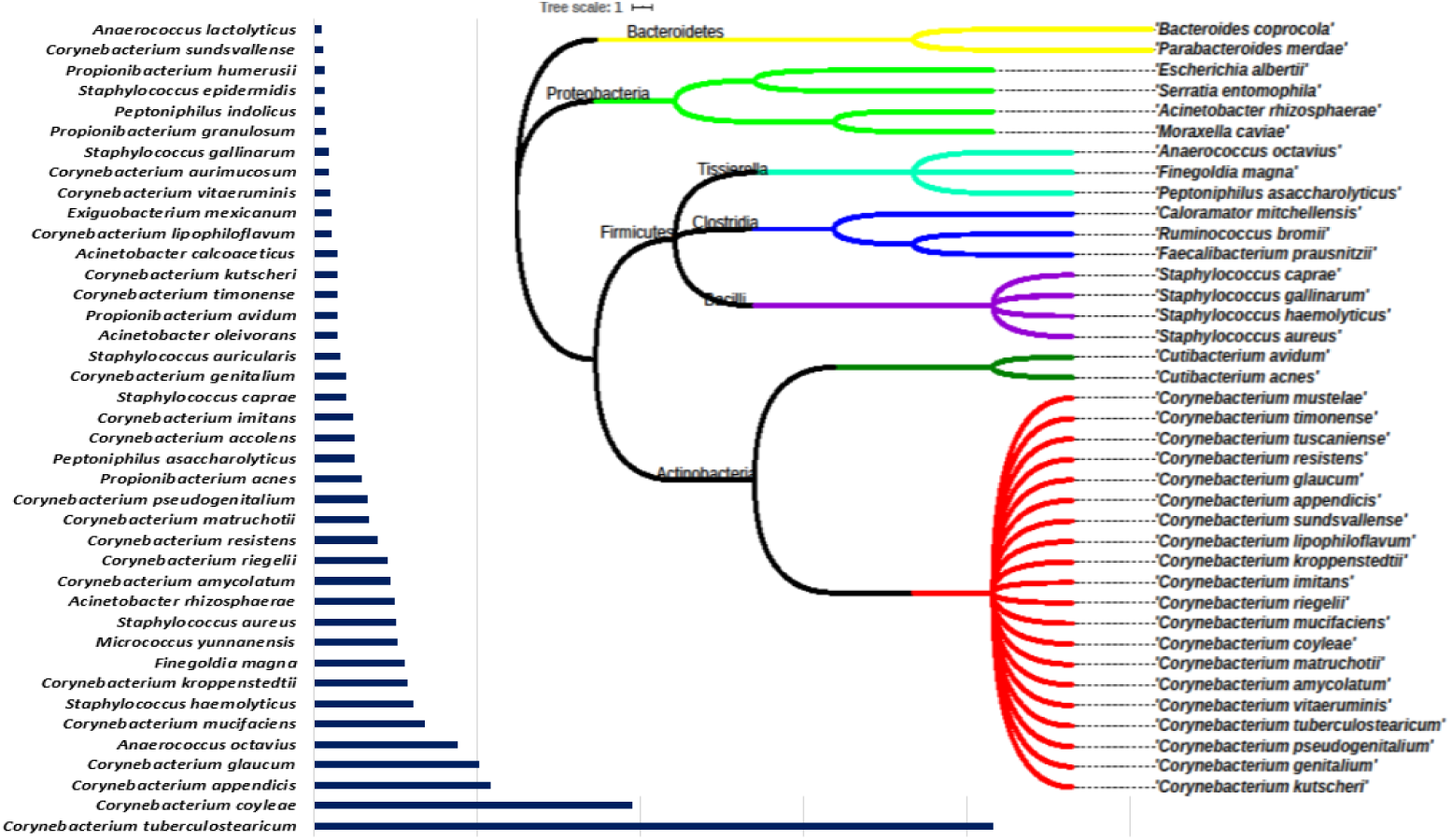
The most relative abundant (%) species in the axilla of female subjects.

**Figure 7:**
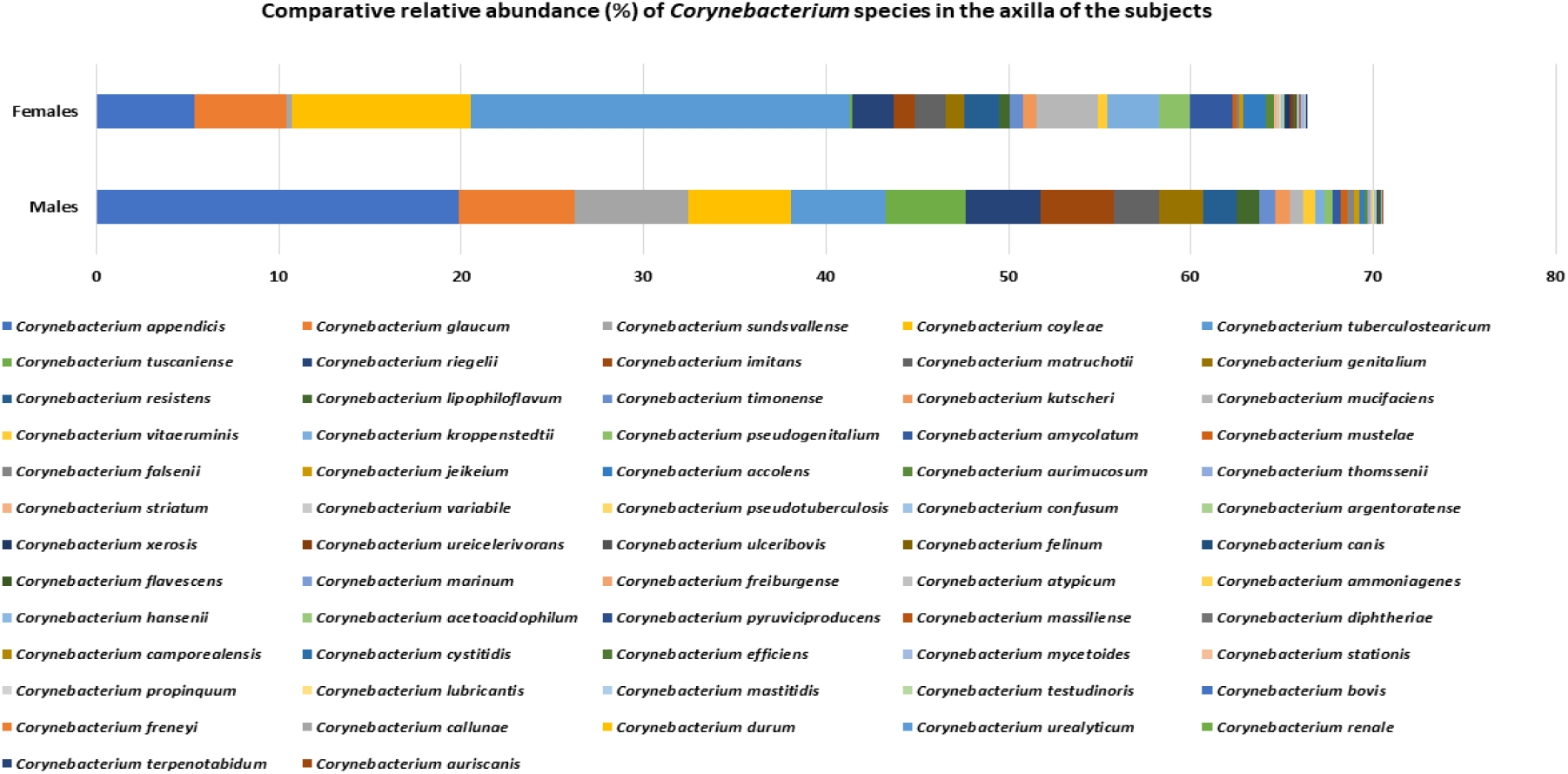
Comparative relative abundance (%) of *Corynebacterium* species in the axilla of the subjects.

Interestingly, 62 Corynebacterium species were found in males, with *Corynebacterium auriscanis* and *Corynebacterium renale* as exclusive, while 63 Corynebacterium species were identified in females with *Corynebacterium casei, Corynebacterium glucuronolyticum* and *Corynebacterium pseudodiphtheriticum* as exclusive.

The axillae of the subjects were also colonized by Lactobacillus species found in 29/35 of female subjects. Among the 29 Lactobacillus species present in female subjects, *Lactobacillus equi, Lactobacillus equicursoris, Lactobacillus plantarum, Lactobacillus fabifermentans, Lactobacillus pantheris* and *Lactobacillus oris* occurred exclusively. The male subjects (31/38) had *Lactobacillus ruminis, Lactobacillus paracasei, Lactobacillus acidifarinae, Lactobacillus casei, Lactobacillus versmoldensis*, and *Lactobacillus hayakitensis* as exclusive (**Figure 8**).

**Figure 8:**
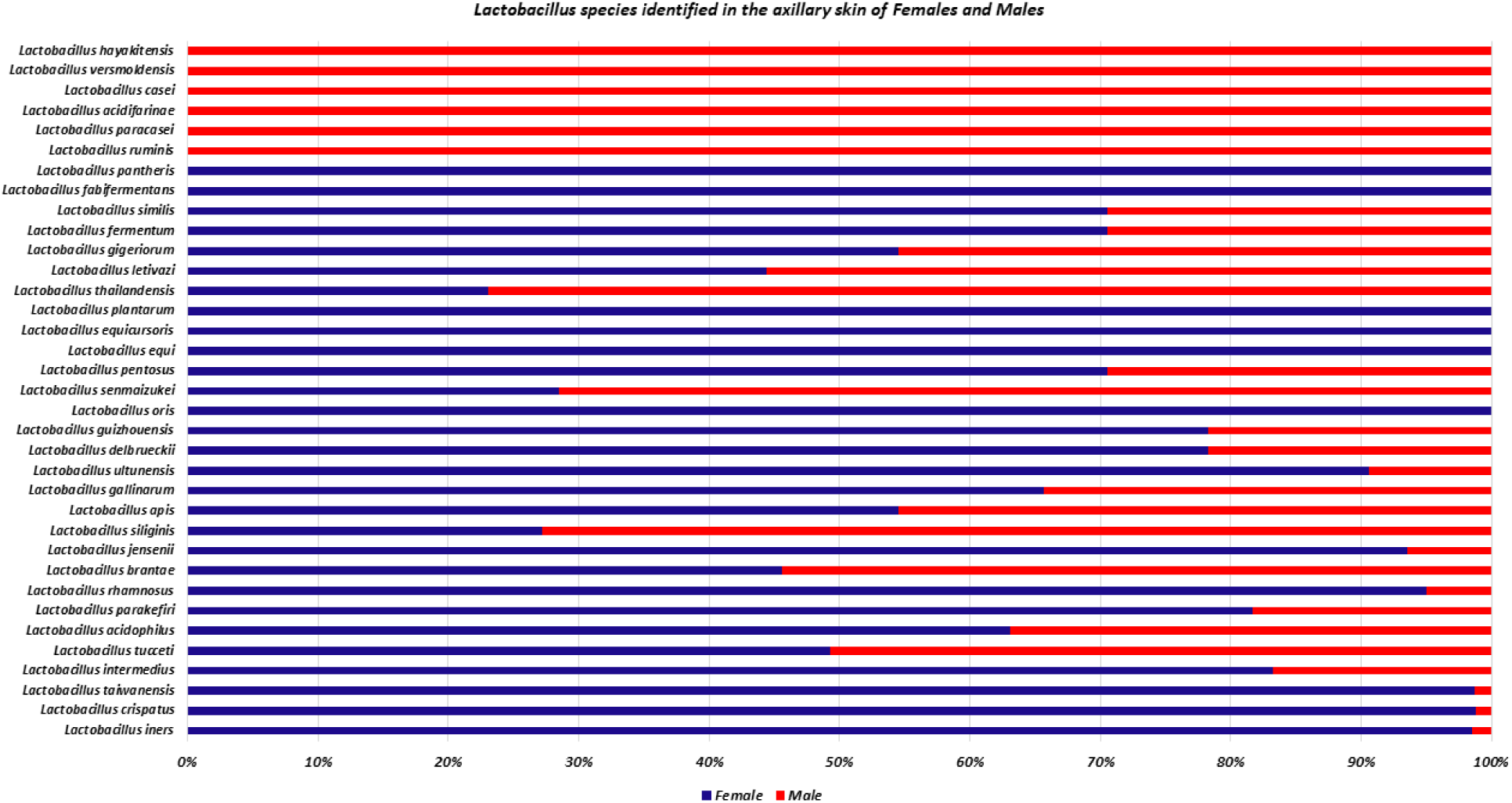
*Lactobacillus* species identified in the axillary skin of female and male subjects.

Staphylococcus species appears to be the second most abundant in both genders, however 34 species were identified in males with *Staphylococcus haemolyticus* (2.82%) as the most abundant species followed by *Staphylococcus aureus* (1.78%), *Staphylococcus gallinarum* (0.80%), *Staphylococcus caprae* (0.73%), *Staphylococcus epidermidis* (0.23%) and others as shown in **Figure 9**.

**Figure 9:**
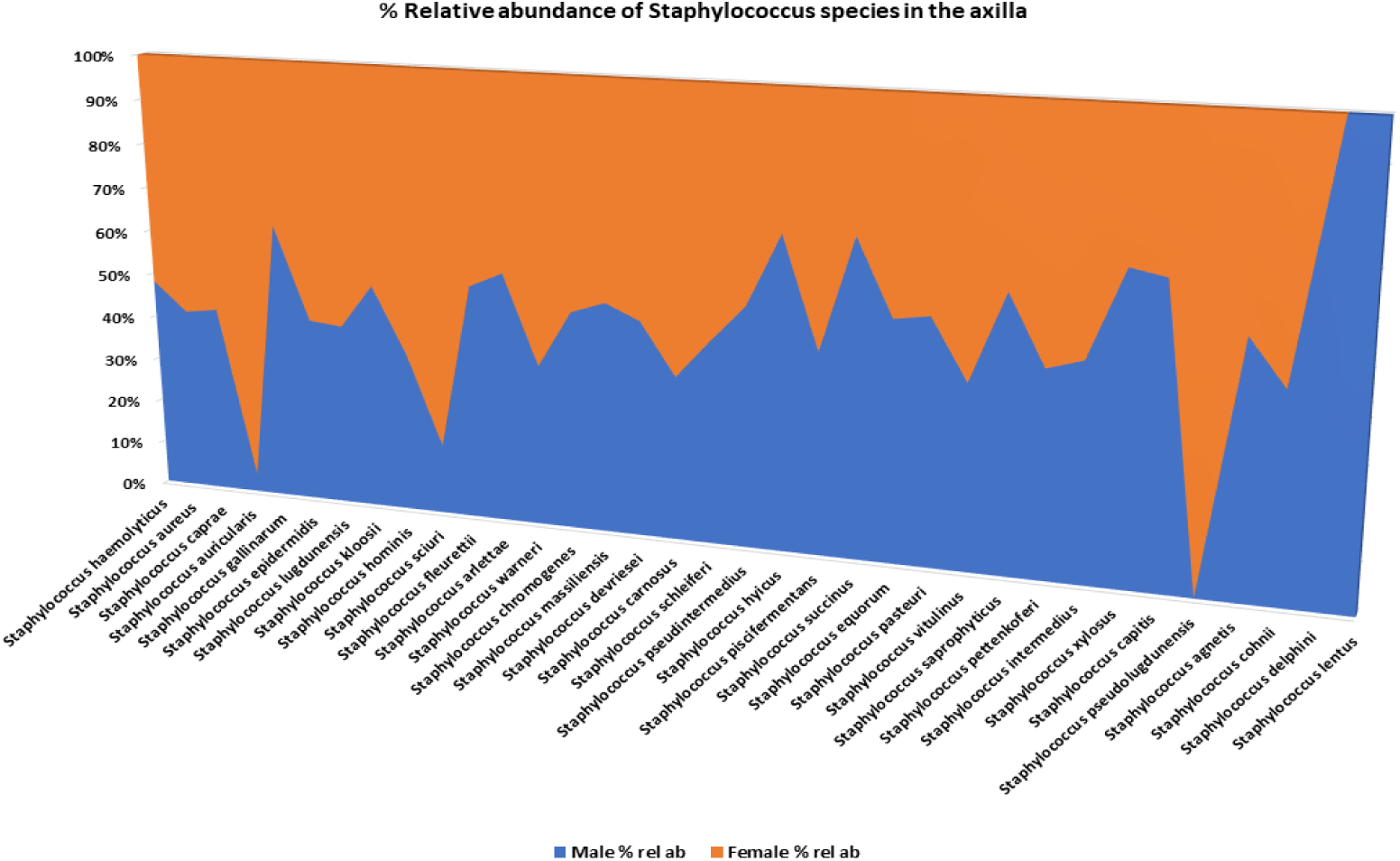
Comparative relative abundance (%) of Staphylococcus species in the axilla of the subjects.

In females, 33 species were found showing *Staphylococcus haemolyticus* (3.04%) as the most relative abundant species followed by *Staphylococcus aureus* (2.51%), *Staphylococcus caprae* (0.99%), *Staphylococcus auricularis* (0.80%), *Staphylococcus gallinarum* (0.46%), *Staphylococcus epidermidis* (0.32%) and others.

The use of antiperspirant/deodorants was reported by 23 males while 15 male subjects stated that they do not use such products. At the genera taxonomic level, there was a significant difference (*P*=0.0075) between those males that reported regular use of antiperspirant/deodorants and those that reported non-use of antiperspirants/deodorants in the relative abundance of Corynebacterium (68.06% vs 42.40%). In contrast, a reverse trend was observed in the relative abundance of Staphylococcus (*P* = 0.047) (2.25% vs 45.10%) as shown in **Figure 10**.

**Figure 10:**
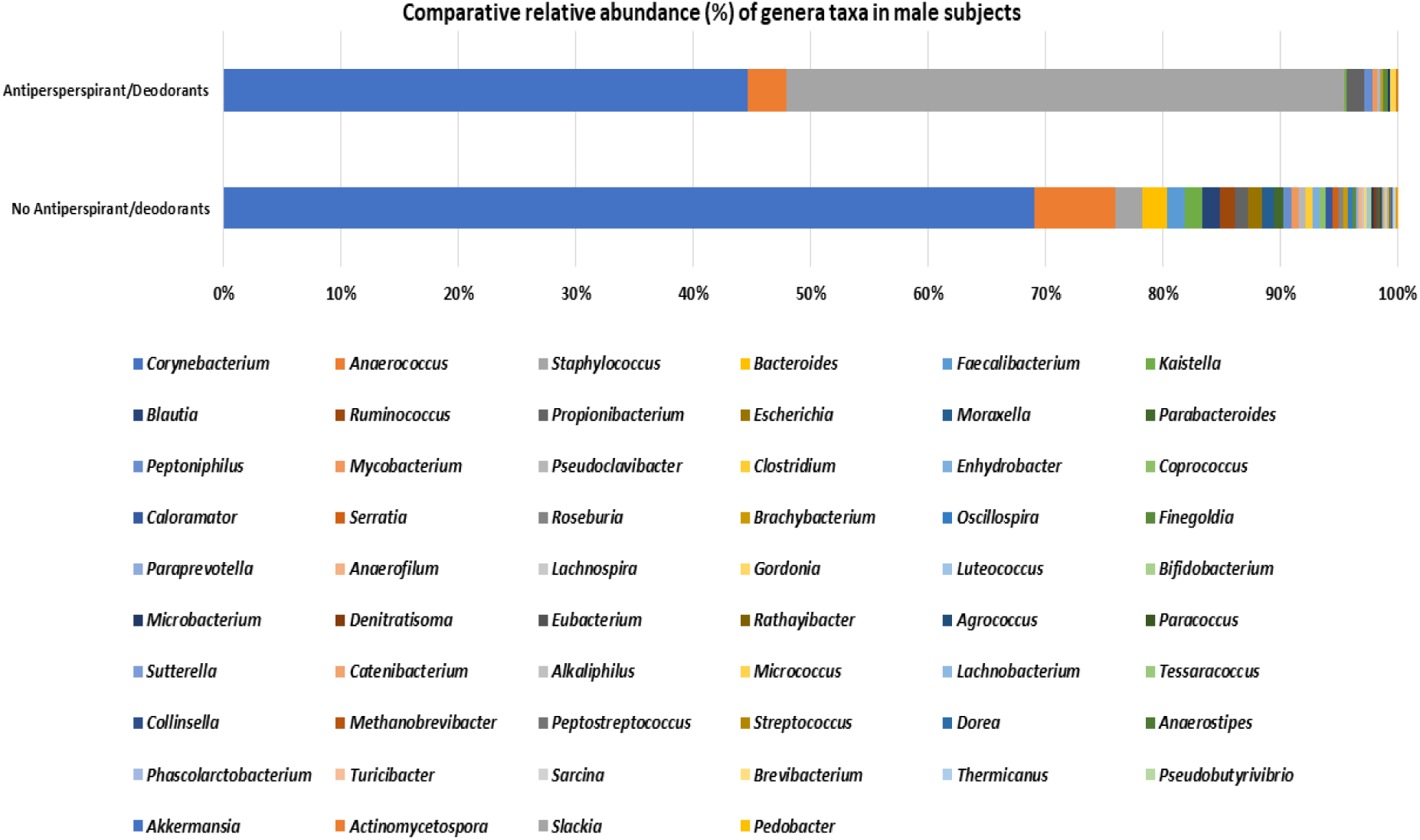
Comparative relative abundance (%) of genera in male subjects.

At the species taxonomic level, males that reported non-use of antiperspirant/deodorants had more relative abundance of *Corynebacterium appendicis* (22.69% vs 14.30%), *Corynebacterium glaucum* (7.43% vs 4.24%), *Corynebacterium tuscaniense* (6.55% vs 0.18%), *Corynebacterium coyleae* (6.27% vs 4.43%), *Corynebacterium imitans* (5.56% vs 0.99%), *Corynebacterium riegelii* (4.88% vs 2.41%) and others represented in **Figure 11**.

**Figure 11:**
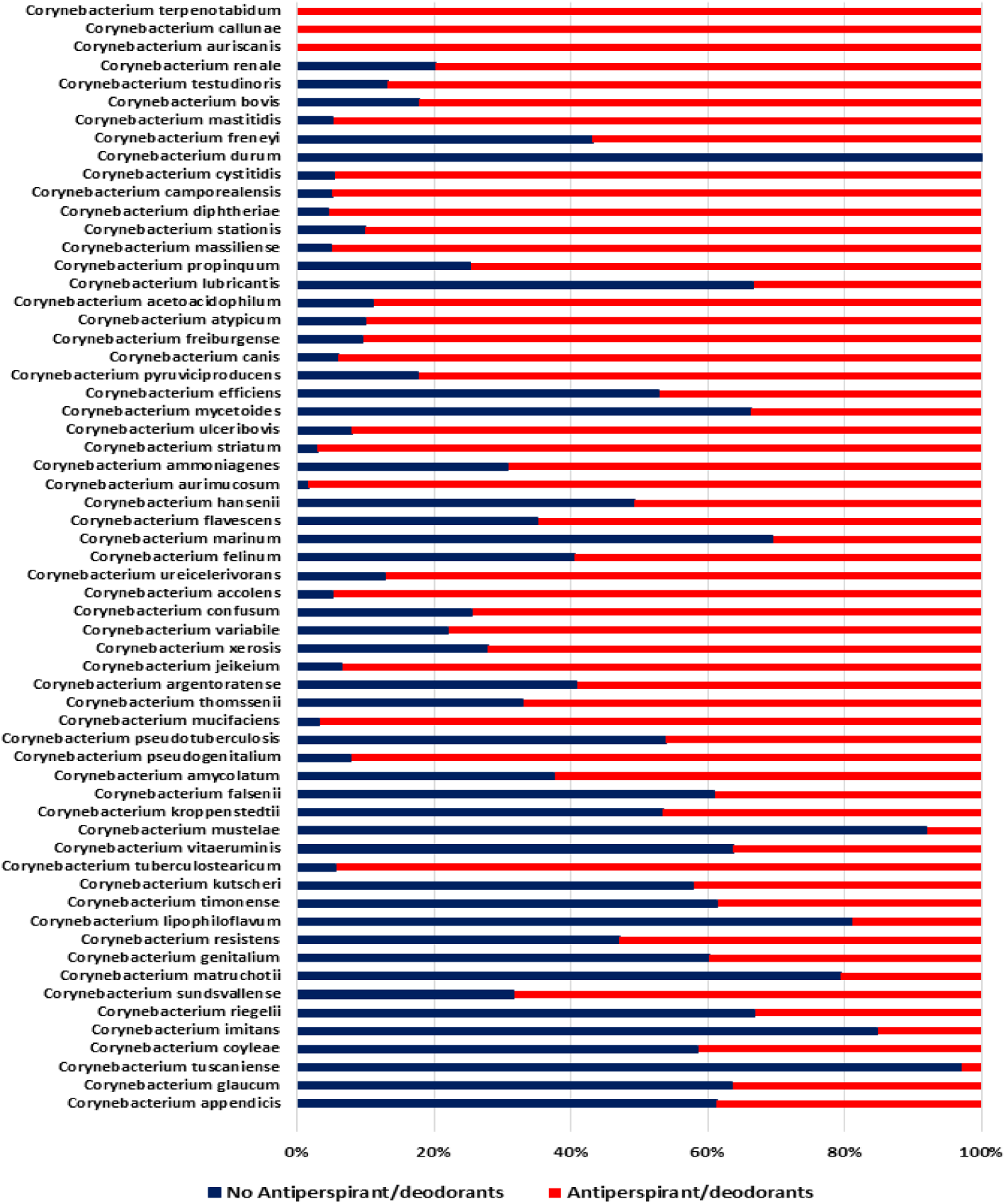
Comparative relative abundance of Corynebacterium species in male subjects that reported regular use of deodorants/antiperspirants and those that don’t.

Conversely, male subjects that reported regular use of antiperspirants/deodorants had more relative abundance of Staphylococcus species than male subjects that reported non-use of antiperspirants/deodorants. For example, *Staphylococcus aureus* (4.76% vs 0.26%), *Staphylococcus gallinarum* (1.85% vs 0.26%), *Staphylococcus haemolyticus* (8.06% vs 0.16%), *Staphylococcus kloosii* (0.13% vs 0.07%), *Staphylococcus caprae* (2.11% vs 0.036%), *Staphylococcus epidermidis* (0.66% vs 0.02%), *Staphylococcus hominis* (0.084% vs 0.003%) and others shown in **Figure 12**.

**Figure 12:**
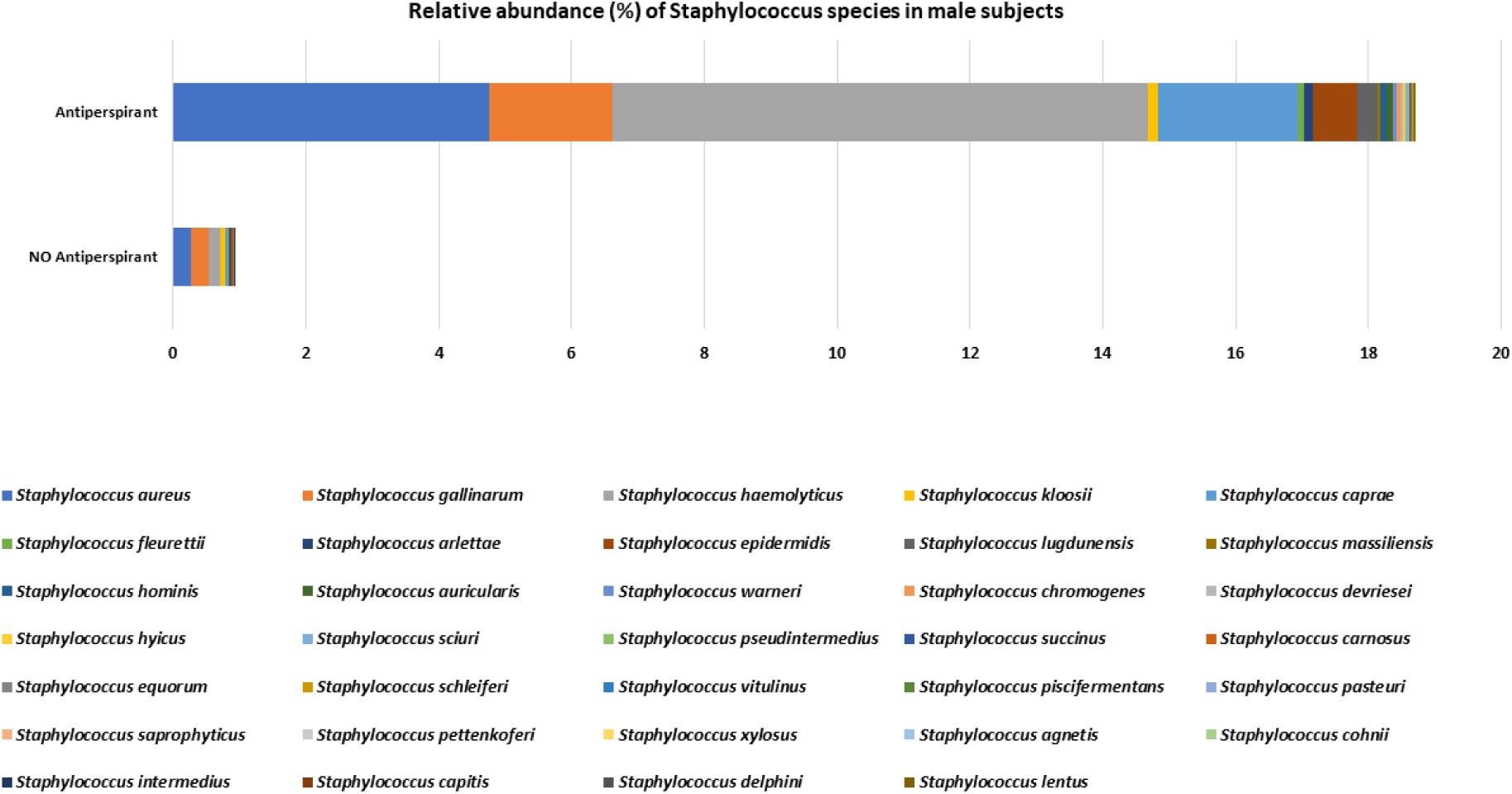
Relative abundance (%) of *Staphylococcus* species in male subjects that reported regular use of deodorants/antiperspirants and those that don’t.

Interestingly, male subjects that reported non-use of antiperspirants/deodorants had more Lactobacillus species in their axilla. Out of 30 Lactobacillus species identified, 23 Lactobacillus were present in non-use of antiperspirants compared with 14 Lactobacillus species found in male subjects that reported regular use of antiperspirants/deodorants.

Although not surprising, out of the 35 female subjects, only 2 reported non-use of antiperspirants/deodorants, while 33 stated regular use of antiperspirants/deodorants. At the genera taxonomic level, *Corynebacterium* (52.26%) was the most relative abundance in the female subjects that reported regular use of antiperspirants/deodorants, followed by *Staphylococcus* (23.10%), *Anaerococcus* (7.25%), *Acinetobacter* (3.11%), *Propionibacterium* (2.04%), *Enhydrobacter* (1.87%), *Micrococcus* (1.83%), *Finegoldia* (1.58%), *Exiguobacterium* (1.15%), *Peptoniphilus* (1.09%). Comparatively, at the species taxonomic level, the relative abundance of the species that occurred 1.0% and above in both male and female subjects that reported regular use of antiperspirants/deodorants is represented in **Figure 13**.

**Figure 13:**
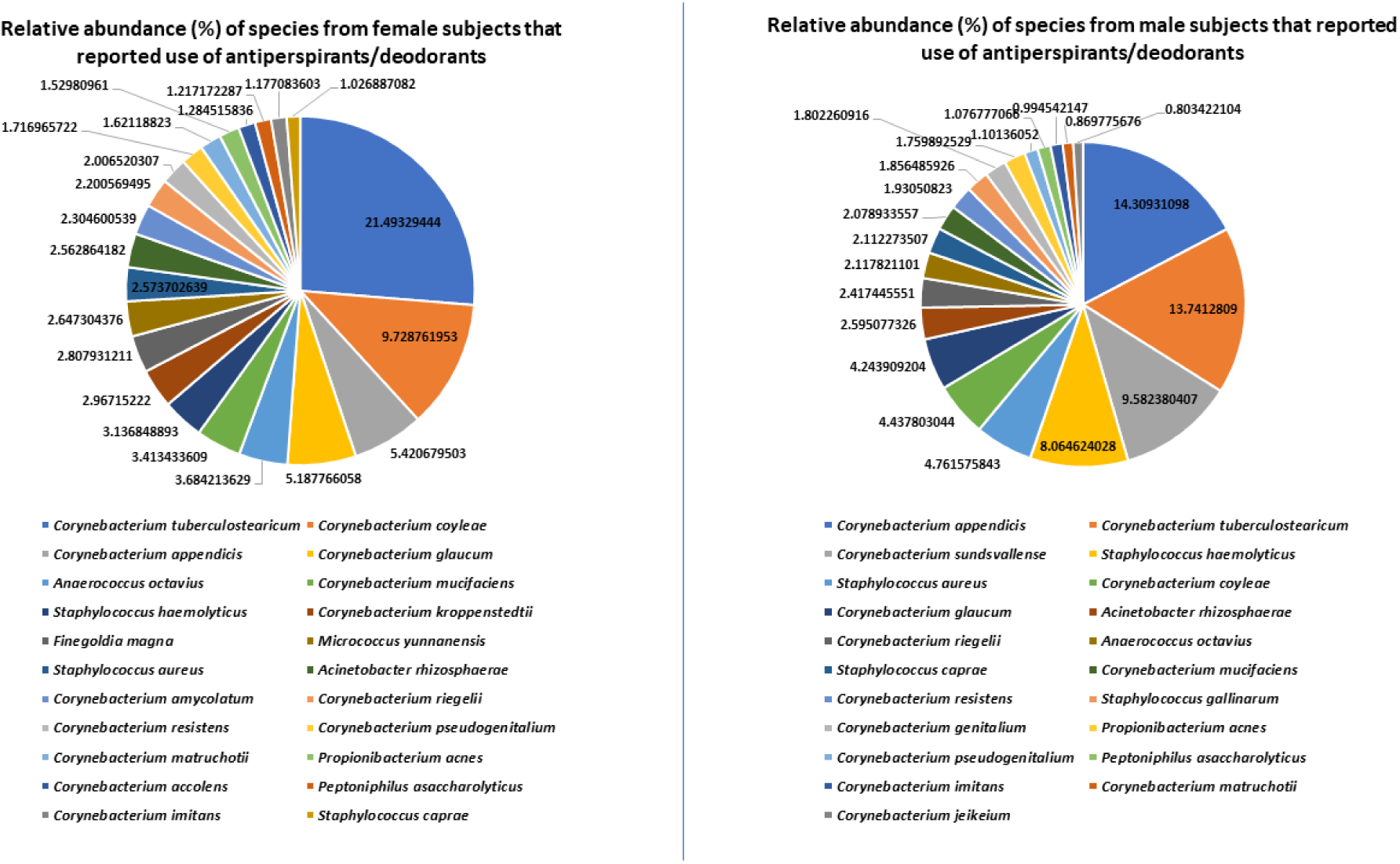
Relative abundance (%) of species from female and male subjects that reported use of antiperspirants/deodorants.

## DISCUSSION

In this study, for the first time in Nigeria, we led an elaborate determination of, and obtained detailed insight into the axillary bacterial communities from both healthy adult male and female students using next generation high throughput sequencing approach. Based on the 16S rRNA dataset obtained, over 99.39% (in males) and 99.92% (in females) of the total sequence reads were assigned to four out of 26 phyla representing *Actinobacteria, Firmicutes Proteobacteria* and *Bacteroidetes*. Our study is in line with the findings of other studies in Europe by Grice et al., (2009a, 2009b) and Costelo et al., (2009). It is noteworthy that out of 747 genera, only three genera *Corynebacterium, Staphylococcus*, and *Anaerococcus* constituted 78.46% of the total reads, which shows that *Corynebacterium* and *Staphylococcus* occupy an importance niche in the human axilla. Similar finding was reported by Callewaert et al (2013).

Our hypothesis appears to be supported by the results obtained at the genera taxonomic level as 74.4% bacterial communities were common to both males and females and 21.8% were exclusively identified in males while 3.8% were exclusive to females. However, it remains to be determined whether the exclusive bacterial communities in males or females confer any differential health benefit, physiological role or pathogenic potential. A relatively higher proportion of *Corynebacterium* was observed in male subjects than females, probably suggesting that gender may play a role. Previous study by Fierer et al (2008) found that *Corynebacterium* tend to colonize male skin especially the hand, more than females. Other studies by Zeeuwen et al (2012) showed that there are differences in the pattern of *Corynebacterium* colonization in the upper buttocks of males and females. It has been postulated that due to anatomical and physiological differences between male and female subjects, especially in hair growth, skin thickness, sex hormones, sweat and sebum production, may be responsible for these microbial differences in the axilla (Giacomoni et al., 2009). In contrast, there are more relative abundance of *Staphylococcus* in females than males, similar to the study conducted on the axillae of adult Belgians (Callewaert et al., 2013). It should be noted that previous studies that utilized culture-dependent methods never reported *Lactobacilli* as being part of the skin and or axillary microbiota. For the fact that in this study, we detected *Lactobacilli* in the axilla of over 82% of females and over 81% of male subjects, though in low relative abundance compared with *Corynebacteria* and *Staphylococci*, indicates that *Lactobacillus* taxa should be considered as part of the normal axillary bacterial community. The female subjects had more relative abundance of *Lactobacillus* taxa than male subjects, which is consistent with the findings of Lebeer et al., (2019) that found a 10-fold higher relative abundance in women than men.

At the species taxonomic level, the proportion of *Corynebacterium appendicis* was more pronounced in males than females in the ratio of 19.86% vs 5.41%, while the reverse was the case for *Corynebacterium tuberculostearicum* found in higher proportion in females than males in the ratio of 20.80% vs 5.18%. The axillary physiological role of this difference remains to be determined, but previous study by Bawdon et al (2015) revealed that *Corynebacterium tuberculostearicum* is a low producer of malodour precursor, a dipeptide-conjugated thioalcohol, particularly S-[1-(2-hydroxyethyl)-1-methylbutyl]-(l)-cysteinylglycine (Cys-Gly-3M3SH). In another study, individuals with higher odour intensities had a greater proportion of *Corynebacterium tuberculostearicum* (Troccaz et al., 2015).

This study revealed that the dominant Staphylococcus species in the sampled population in both male and female subjects were *Staphylococcus haemolyticus* and *Staphylococcus aureus*. This is inconsistent with the study by Egert et al. (2011) that showed the dominant *Staphylococcus* species in the axilla were *Staphylococcus epidermidis* and *Staphylococcus hominis*.

By clustering the bacterial communities from males that reported regular use of antiperspirant/deodorants and those that reported non-use of antiperspirants/deodorants, we observed that the relative abundance of *Corynebacterium* (68.06% vs 42.40%) was higher in those reported regular use of deodorants. The implication of this is that the use of deodorants/antiperspirants facilitates the proliferation of *Corynebacterium* species as high levels of strong body odour were observed by Taylor et al (2003) in individuals with a microbiota dominated by *Corynebacterium*. Interestingly, in this study, those that reported non-use of deodorants/antiperspirants, the axillae were dominated by *Staphylococci*, as *staphylococci*-dominated axillae revealed low levels of odour (Taylor et al 2003). We found out a significant reduction in the species richness and diversities of *Lactobacillus* taxa from those that reported regular use of deodorants/antiperspirants than those that do not, thus suggesting that beneficial bacteria such as Lactobacilli are impacted negatively by the use of these products.

In this study, *Staphylococcus haemolyticus* (8.06% vs 0.16%), and *Staphylococcus hominis* (0.084% vs 0.003%) were more in male subjects that reported regular use of deodorants/antiperspirants, which suggests that they may be having more malodour as *Staphylococcus hominis* and *Staphylococcus haemolyticus* have been identified as efficient bio-transformers of Cys-Gly-3M3SH (Bawdon et al., 2015).

The limitations associated with this study especially on the use of antiperspirants/deodorants verges on the inability to collect information on the exact regular products used. The subjects that used deodorants and or antiperspirants may have different levels of bacterial communities.

## CONCLUSION

We have shown in this study that the axilla of the sampled students is composed of bacterial communities that largely represented *Actinobacteria, Firmicutes Proteobacteria* and *Bacteroidetes*. A relatively higher proportion of *Corynebacterium* was observed in male subjects than females, while more relative abundance of *Staphylococcus* was found in females than males. This study detected *Lactobacilli* in the axilla of over 82% of female and over 81% of male subjects, though in low relative abundance which suggests that *Lactobacillus* taxa might be considered as part of the normal axillary bacterial community. The study also revealed that the relative abundance of *Corynebacterium* (68.06% vs 42.40%) was higher in those reported regular use of deodorants. The implication of this is that the use of deodorants/antiperspirants may facilitate the proliferation of malodour-producing *Corynebacterium* and *Staphylococcus* species, while decreasing beneficial bacteria such as *Lactobacilli* in the axilla.

## Supplementary Tables

**Supplementary Table 1**: Bacterial species exclusively identified in male subjects

**Supplementary Table 2**: Bacterial species exclusively identified in the axilla of female subjects

## AUTHOR CONTRIBUTIONS

KCA and NRA conceived and designed the study. KCA sourced for funding, wrote the protocol, did literature search, did bioinformatics analysis, interpreted the data and wrote the final manuscript. VM did the survey experiments, collected the samples, did initial literature searches, taxonomic data organization & statistical analysis, wrote the draft manuscript and recruited the subjects. KCA and NRA supervised the study and approved the submitted manuscript.

## ACKNOWLEDGMENTS

We sincerely thank uBiome Inc, San Francisco, California, USA (uBiome has been liquidated and bought over by a Korean company) for awarding a grant-in-kind to Dr. Kingsley Anukam and for carrying out the metagenomics sequencing. We gratefully acknowledge the student volunteers who freely participated in the study. KCA is a visiting reader to the Departments of Medical Laboratory Science and Pharmaceutical Microbiology, Nnamdi Azikiwe University.

