## Supplemental Table 1 for "Axillary Microbiota Compositions from Men and Women in a Tertiary Institution-South East Nigeria: Effects of Deodorants/Antiperspirants on Bacterial Communities"

**Supplementary Table 1: Bacterial species exclusively identified in male subjects**

| | | | |
|---|---|---|---|
| Acholeplasma ales | Clostridium magnum | Lactobacillus ruminis | Pseudomonas collierea |
| Acholeplasma cavigenitalium | Clostridium malenominatum | Lactobacillus salivarius | Pseudomonas jinjuensis |
| Acholeplasma equifetale | Clostridium nitrophenolicum | Lactobacillus versmoldensis | Pseudomonas luteola |
| Acholeplasma granularum | Clostridium paraputrificum | Lactococcus lactis | Pseudomonas panipatensis |
| Acholeplasma hippikon | Clostridium perfringens | Legionella fallonii | Pseudomonas poae |
| Acidaminobacter hydrogenoformans | Clostridium proteolyticum | Legionella taurinensis | Pseudomonas tropicalis |
| Acidaminococcus intestini | Clostridium proteolyticus | Legionella worsleiensis | Pseudonocardia acaciae |
| Acidiphilium acidophilum | Clostridium saccharoperbutylacetonicum | Lentzea californiensis | Pseudonocardia asaccharolytica |
| Acidiphilium organovorum | Clostridium sulfidigenes | Leptotrichia buccalis | Pseudonocardia hydrocarbonoxydans |
| Acidiphilium symbioticum | Clostridium termitidis | Leptotrichia shahii | Pseudoxanthomonas mexicana |
| Acidisoma tundrae | Clostridium tetani | Leptotrichia wadei | Pullulanibacillus naganoensis |
| Acidovorax facilis | Clostridium thermoalcaliphilum | Leucobacter albus | Rhizobium mesoamericanum |
| Acinetobacter indicus | Clostridium tunisiense | Lewinella marina | Rhodanobacter thiooxydans |
| Acinetobacter venetianus | Cohnella fontinalis | Luteimonas aquatica | Rhodobacter blasticus |
| Actinoallomurus luridus | Cohnella laeviribosi | Luteimonas mephitis | Rhodobacter gluconicum |
| Actinobacillus pleuropneumoniae | Collinsella stercoris | Lysobacter daejeonensis | Rhodobium gokarnense |
| Actinobaculum urinale | Coprobacillus cateniformis | Lysobacter enzymogenes | Rhodococcus fascians |
| Actinokineospora diospyrosa | Corallococcus exiguus | Lysobacter yangpyeongensis | Rhodococcus zopfii |
| Actinomadura cremea | Coriobacterium glomerans | Marichromatium gracile | Rhodovulum imhoffii |
| Actinomadura macra | Corynebacterium auriscanis | Marinobacter santoriniensis | Rhodovulum iodosum |
| Actinomyces georgiae | Corynebacterium renale | Marinobacterium stanieri | Rhodovulum robiginosum |
| Actinomyces suimastitidis | Crocosphaera watsonii | Marinococcus salidurans | Rickettsia hulinii |
| Actinoplanes auranticolor | Cupriavidus gilardii | Marinomonas pontica | Rickettsia marmionii |
| Actinoplanes garbadinensis | Cupriavidus pauculus | Meiothermus granaticius | Rickettsia monacensis |
| Actinopolyspora xinjiangensis | Curtobacterium herbarum | Mesonia algae | Riemerella columbina |
| Aequorivita crocea | Cyanobacterium aponinum | Mesorhizobium camelthorni | Rikenella microfusus |

Aerococcus sanguinicola
Agrobacterium undicola
Agromyces allii
Agromyces fucosus
Agromyces mediolanus
Alkalibacillus haloalkaliphilus
Alkalibacterium subtropicum
Alkaliphilus metalliredigens
Alkaliphilus peptidifermentans
Allochromatium palmeri
Aminobacter aganoensis
Aminobacter ciceronei
Aminobacterium colombiense
Anaerofustis stercorihominis
Anaerospora hongkongensis
Anaerovibrio lipolyticus
Ancylobacter aquaticus
Aneurinibacillus danicus
Aquimarina macrocephali
Arcanobacterium bernardiae
Arthrobacter crystallopoietes
Arthrobacter kerguelensis
Azohydromonas australica
Azospira restricta
Azospirillum palatum
Azospirillum zeae
Bacillus aerophilus
Bacillus arbutinivorans

Cyanobacterium stanieri
Cycloclasticus oligotrophus
Deinococcus aeria
Deinococcus aerius
Deinococcus alpinitundrae
Deinococcus deserti
Deinococcus geothermalis
Deinococcus gobiensis
Deinococcus murrayi
Deinococcus reticulitermitis
Deinococcus yunweiensis
Demequina salsinemoris
Dermatophilus congolensis
Desulfacinum subterraneum
Desulfitobacterium chlororespirans
Desulfofrigus fragile
Desulfofrigus oceanense
Desulfonatronovibrio
Desulfonatronovibrio hydrogenovorans
Desulfosporosinus lacus
Desulfotomaculum australicum
Desulfovibrio butyratiphilus
Desulfovibrio caledoniensis
Desulfovibrio carbinolicus
Desulfovibrio psychrotolerans
Desulfurispirillum indicum
Desulfuromonas thiophila
Devosia chinhatensis

Mesorhizobium opportunistum
Mesorhizobium septentrionale
Methanobrevibacter acididurans
Methanobrevibacter woesei
Methylobacillus flagellatus
Methylobacterium adhaesivum
Methylobacterium dankookense
Methylobacterium mesophilicum
Methylobacterium organophilum
Methylocella silvestris
Methylophaga aminisulfidivorans
Microbacterium aoyamense
Microbacterium flavum
Microbacterium halotolerans
Microbacterium koreense
Microbacterium terricola
Microbacterium thalassium
Microbacterium xinjiangensis
Micrococcus endophyticus
Micromonospora aquatica
Micromonospora fulvoviolacea
Mobiluncus mulieris
Mogibacterium vescum
Moraxella equi
Morganella psychrotolerans
Muricauda lutimaris
Mycobacterium acapulcensis
Mycobacterium brasiliensis

Rivularia atra
Roseateles depolymerans
Roseococcus suduntuyensis
Roseomonas lacus
Roseospira visakhapatnamensis
Rothia dentocariosa
Rubellimicrobium mesophilum
Rubellimicrobium roseum
Rubrobacter xylanophilus
Ruminobacter amylophilus
Ruminococcus callidus
Saccharospirillum impatiens
Salisaeta longa
Sarcina ventriculi
Scardovia inopinata
Schlegelella aquatica
Scytonema hofmanni
Sedimentibacter saalensis
Segetibacter aerophilus
Serratia nematodiphila
Sharpea azabuensis
Shewanella amazonensis
Shewanella decolorationis
Shewanella upenei
Shewanella vesiculosa
Shimazuella kribbensis
Shinella granuli
Sinorhizobium fredii

| | | | |
|---|---|---|---|
| Bacillus butanolivorans | Devosia ginsengisoli | Mycobacterium fuerthensis | Slackia heliotrinireducens |
| Bacillus cereus | Devosia hwasunensis | Mycobacterium noviomagense | Slackia piriformis |
| Bacillus coagulans | Devosia limi | Mycobacterium obuense | Sneathia sanguinegens |
| Bacillus firmus | Devosia riboflavina | Mycoplasma auris | Snowella rosea |
| Bacillus flexus | Dickeya dianthicola | Mycoplasma coccoides | Sodalis glossinidius |
| Bacillus gibsonii | Dietzia papillomatosis | Mycoplasma edwardii | Sphingobacterium thalpophilum |
| Bacillus halodurans | Dokdonella fugitiva | Myroides profundi | Sphingobium faniae |
| Bacillus horikoshii | Dolichospermum curvum | Myxococcus xanthus | Sphingobium olei |
| Bacillus longiquaesitum | Dyadobacter alkalitolerans | Natronincola ferrireducens | Sphingomonas abaci |
| Bacillus marisflavi | Ectothiorhodospira haloalkaliphila | Neisseria cinerea | Sphingomonas asaccharolytica |
| Bacillus nealsonii | Ectothiorhodospira imhoffii | Neisseria flavescens | Sphingomonas fennica |
| Bacillus niabensis | Edwardsiella hoshinae | Niabella aurantiaca | Sphingomonas hankookensis |
| Bacillus olivae | Eggerthella lenta | Niabella soli | Sphingomonas hunanensis |
| Bacillus oryzae | Eikenella corrodens | Niastella koreensis | Sphingomonas japonica |
| Bacillus plakortidis | Emticicia ginsengisoli | Nisaea nitritireducens | Sphingomonas phyllosphaerae |
| Bacillus pseudofirmus | Emticicia oligotrophica | Nitrosococcus watsoni | Sphingomonas pruni |
| Bacillus psychrosaccharolyticus | Enterobacter aceae | Nitrospira moscoviensis | Sphingomonas roseiflava |
| Bacillus thioparans | Enterobacter cowanii | Nocardia araoensis | Sphingopyxis chilensis |
| Bacteroides barnesiae | Enterobacter ludwigii | Nocardia harenae | Sphingopyxis ginsengisoli |
| Bacteroides chinchillae | Enterobacter nickellidurans | Nocardia higoensis | Sphingopyxis granuli |
| Bacteroides coprophilus | Enterococcus columbae | Nocardia jinanensis | Sphingopyxis panaciterrae |
| Bacteroides eggerthii | Enterococcus italicus | Nocardia novocastrensa | Sphingopyxis witflariensis |
| Bacteroides gallinarum | Enterococcus rottae | Nocardioides islandensis | Spirosoma rigui |
| Bacteroides intestinalis | Erwinia billingiae | Novosphingobium subterraneum | Sporosarcina ureae |
| Bacteroides salanitronis | Erwinia oleae | Oceanimonas smirnovii | Staphylococcus delphini |
| Bacteroides salyersiae | Erwinia pyrifoliae | Oceanospirillum multiglobuliferum | Staphylococcus lentus |
| Balneola vulgaris | Erwinia rhapontici | Ochrobactrum anthropi | Stenotrophomonas nitritireducens |
| Bdellovibrio bacteriovorus | Erythrobacter flavus | Ochrobactrum intermedium | Steroidobacter denitrificans |
| Bdellovibrio exovorus | Erythromicrobium ramosum | Olivibacter ginsengisoli | Streptococcus dentapri |

Bellilinea caldifistulae
Bifidobacterium bombi
Bifidobacterium boum
Bifidobacterium kashiwanohense
Bifidobacterium magnum
Bifidobacterium scardovii
Blastococcus aggregatus
Brevibacillus ginsengisoli
Bulleidia extructa
Bulleidia moorei
Burkholderia lata
Burkholderia multivorans
Burkholderia phenoliruptrix
Burkholderia seminalis
Burkholderia vietnamiensis
Butyricimonas synergistica
Butyrivibrio hungatei
Caldanaerobacter hydrothermalis
Caldicellulosiruptor bescii
Caloramator australicus
Caloramator uzoniensis
Caloramator viterbiensis
Campylobacter concisus
Candidatus Brodae
Candidatus Contubernalis Alkalaceticum
Candidatus Endobugula
Candidatus Glomeribacter
Candidatus Methylacidiphilum
Candidatus Nitrososphaera
Candidatus Phlomobacter Fragariae

Eubacterium callanderi
Eubacterium cylindroides
Facklamia hominis
Ferrimonas kyonanensis
Fibrobacter intestinalis
Filifactor alocis
Filifactor villosus
Frankia alni
Friedmanniella okinawensis
Frigoribacterium faeni
Fructobacillus fructosus
Fusibacter paucivorans
Fusobacterium canifelinum
Fusobacterium simiae
Gallionella ferruginea
Gemella bergeri
Gemmatimonas aurantiaca
Geobacillus thermoglucosidans
Geobacter grbiciae
Geobacter pelophilus
Geobacter pickeringii
Georgenia halophila
Gillisia limnaea
Gillisia sandarakina
Glycomyces tenuis
Gordonia aichiensis
Gordonia defluvii
Gordonia desulfuricans
Gordonia westfalica
Haemophilus quentini

Olivibacter soli
Oxalobacter formigenes
Paenibacillus amylolyticus
Paenibacillus barcinonensis
Paenibacillus chitinolyticus
Paenibacillus donghaensis
Paenibacillus ehimensis
Paenibacillus filicis
Paenibacillus forsythiae
Paenibacillus ginsengagri
Paenibacillus humicus
Paenibacillus illinoisensis
Paenibacillus mendelii
Paenibacillus pasadenensis
Paenibacillus phyllosphaerae
Paenibacillus thailandensis
Paenibacillus woosongensis
Parabacteroides gordonii
Paracoccus alcaliphilus
Paracoccus halotolerans
Paracoccus kamogawaensis
Paracoccus kocurii
Paracoccus solventivorans
Paracoccus thiocyanatus
Parapedobacter koreensis
Parvibaculum lavamentivorans
Pasteurella pneumotropica
Pediococcus argentinicus
Pediococcus pentosaceus
Pediococcus stilesii

Streptococcus intermedius
Streptococcus marimammalium
Streptococcus massiliensis
Streptococcus minor
Streptococcus parasanguinis
Streptococcus phocae
Streptococcus ursoris
Streptomonospora halophila
Streptomyces aculeolatus
Streptomyces beijiangensis
Streptomyces espinosus
Streptomyces eurythermus
Streptomyces griseoaurantiacus
Streptomyces noursei
Streptomyces olivogriseus
Streptomyces panayensis
Streptomyces pulcher
Streptomyces roseogilvus
Streptomyces roseosporus
Streptomyces sannanensis
Streptomyces scopiformis
Streptomyces tacrolimicus
Streptomyces tendae
Streptomyces thermoluteus
Streptomyces vastus
Streptomyces viridis
Streptomyces zinciresistens
Streptosporangium yunnanense
Succinivibrio dextrinosolvens
Sutterella parvirubra

Candidatus Phytoplasma Prunorum
Candidatus Portiera
Candidatus Protochlamydia Amoebophila
Candidatus Rhabdochlamydia
Candidatus Solibacter
Candidatus Tammella Caduceiae
Candidatus Xiphinematobacter
Capnocytophaga gingivalis
Carnobacterium gallinarum
Catenibacterium mitsuokai
Catenulispora rubra
Cellulomonas flavigena
Cellvibrio ostraviensis
Chlorobaculum limnaeum
Chromobacterium piscinae
Chromobacterium subtsugae
Chryseobacterium hispanicum
Chryseobacterium wanjuense
Chthoniobacter flavus
Citrobacter braakii
Citrobacter freundii
Citrobacter werkmanii
Clostridium acidisoli
Clostridium aldrichii
Clostridium aurantibutyricum
Clostridium baratii
Clostridium bovipellis
Clostridium butyricum
Clostridium cavendishii
Clostridium cellulovorans
Haererehalobacter ostenderis
Hahella antarctica
Haliangium ochraceum
Halochromatium salexigens
Halococcus dombrowskii
Halococcus saccharolyticus
Halococcus thailandensis
Halomonas almeriensis
Halomonas eurihalina
Halomonas fontilapidosi
Halomonas glaciei
Halomonas nitroreducens
Halomonas sabkhae
Halomonas venusta
Halorubrum jeotgali
Haloterrigena limicola
Heliorestis baculata
Herbaspirillum frisingense
Herbaspirillum seropedicae
Hydrogenophaga intermedia
Hymenobacter gelipurpurascens
Hymenobacter ocellatus
Hymenobacter rigui
Hyphomicrobium aestuarii
Hyphomicrobium hollandicum
Hyphomicrobium vulgare
Janibacter corallicola
Janibacter terrae
Jannaschia seohaensis
Jiangella alkaliphila
Pedobacter aquatilis
Pedobacter ginsengisoli
Pedobacter kribbensis
Pedobacter kwangyangensis
Pedobacter rhizospharae
Pedobacter westerhofensis
Pedomicrobium manganicum
Phaeobacter arcticus
Photobacterium lutimaris
Phyllobacterium bourgognense
Pirellula staleyi
Planctomyces brasiliensis
Planctomyces limnophilus
Planctomyces maris
Planktothricoides raciborskii
Planococcus rifietoensis
Planomicrobium flavidum
Polaromonas jejuensis
Porphyromonas endodontalis
Prevotella albensis
Prevotella histicola
Prevotella intermedia
Prevotella loescheii
Prevotella nanceiensis
Prevotella nigrescens
Prevotella oris
Prevotella oulorum
Prevotella shahii
Prevotella veroralis
Promicromonospora sukumoe
Sutterella sanguinus
Symbiobacterium toebii
Tepidimonas arfidensis
Tepidimonas thermarum
Terribacillus goriensis
Terribacillus halophilus
Tetragenococcus halophilus
Tetrasphaera vanveenii
Thauera chlorobenzoica
Thauera mechernichensis
Thauera terpenica
Thermobaculum terrenum
Thermodesulfovibrio thiophilus
Thermomonas haemolytica
Thioalkalimicrobium aerophilum
Thiobacillus thiophilus
Thiohalorhabdus denitrificans
Thiorhodococcus mannitoliphagus
Thiorhodococcus pfennigii
Thiothrix caldifontis
Thiothrix fructosivorans
Thiothrix lacustris
Thiothrix nivea
Trabulsiella farmeri
Trabulsiella guamensis
Trabulsiella odontotermitis
Treponema paraluiscuniculi
Treponema porcinum
Treponema socranskii
Treponema succinifaciens

Clostridium chartatabidum
Clostridium chromoreductans
Clostridium fallax
Clostridium frigidicarnis
Clostridium gasigenes
Clostridium hiranonis
Clostridium homopropionicum
Clostridium homopropionicum
Clostridium kluyveri
Clostridium ljungdahlii
Clostridium lundense

Kineosporia babensis
Kitasatospora setae
Kocuria atrinae
Kocuria halotolerans
Kosmotoga arenicorallina
Kouleothrix aurantiaca
Kushneria indalinina
Lactobacillus acidifarinae
Lactobacillus casei
Lactobacillus hayakitensis
Lactobacillus paracasei

Propionigenium modestum
Prosthecobacter debontii
Prosthecobacter fluviatilis
Proteus hauseri
Proteus penneri
Pseudaminobacter salicylatoxidans
Pseudoclavibacter helvolus
Pseudomonas agarici
Pseudomonas alcaligenes
Pseudomonas azotoformans
Pseudomonas benzenivorans

Vagococcus penaei
Veillonella denticariosi
Vibrio gazogenes
Vibrio mytili
Virgibacillus byunsanensis
Virgibacillus olivae
Williamsia marianensis
Wolbachia pipientis
