## Supplemental Table 2 for "Axillary Microbiota Compositions from Men and Women in a Tertiary Institution-South East Nigeria: Effects of Deodorants/Antiperspirants on Bacterial Communities"

**Supplementary Table 2: Bacterial species exclusively identified in the axilla of female subjects**

| | | | |
|---|---|---|---|
| Acetobacter indonesiensis | Cryocola antiquus | Pediococcus acidilactici | Shewanella morhuae |
| Acetobacter orleanensis | Dactylosporangium fulvum | Pediococcus cellicola | Shewanella oneidensis |
| Acinetobacter bouvetii | Deinococcus grandis | Pediococcus siamensis | Shewanella putrefaciens |
| Acinetobacter lwoffii | Deinococcus hopiensis | Photobacterium aquimaris | Slackia exigua |
| Actinoalloteichus nanshanensis | Deinococcus proteolyticus | Photobacterium ganghwense | Solibacillus silvestris |
| Actinobaculum schaalii | Deinococcus yavapaiensis | Photobacterium kishitanii | Sphingobacterium faecium |
| Actinomyces cardiffensis | Desulfotalea arctica | Photobacterium rosenbergii | Sporosarcina soli |
| Actinomyces europaeus | Desulfovibrio aceae | Planomicrobium chinense | Staphylococcus pseudolugdunensis |
| Actinomyces hyovaginalis | Dietzia schimae | Pontibacillus chungwhensis | Stenotrophomonas chelatiphaga |
| Actinomyces neuii | Elizabethkingia anophelis | Pontibacter korlensis | Sterolibacterium denitrificans |
| Agrococcus jejuensis | Enterococcus faecium | Pontibacter xinjiangensis | Streptacidiphilus jiangxiensis |
| Alishewanella fetalis | Enterococcus gallinarum | Prevotella disiens | Streptococcus agalactiae |
| Alkalibacterium psychrotolerans | Exiguobacterium profundum | Prevotella salivae | Streptococcus ictaluri |
| Amycolatopsis halophila | Exiguobacterium taiwanense | Prevotella stercorea | Streptococcus orisratti |
| Amycolatopsis mediterranei | Ferrimicrobium acidiphilum | Prevotella tannerae | Streptococcus urinalis |
| Arcobacter cryaerophilus | Flavobacterium anhuiense | Providencia alcalifaciens | Streptomyces luteireticuli |
| Arthrobacter psychrolactophilus | Flavobacterium kamogawaensis | Providencia burhodogranariea | Streptomyces nanchangensis |
| Atopobium minutum | Flavobacterium suncheonense | Pseudoalteromonas gracilis | Streptomyces poonensis |
| Azomonas insignis | Flavobacterium weaverense | Pseudoalteromonas haloplanktis | Streptomyces radiopugnans |
| Azospirillum canadense | Geobacillus caldoxylosilyticus | Pseudochrobactrum saccharolyticum | Streptomyces variegatus |
| Bacillus alcaliinulinus | Geobacillus gargensis | Pseudomonas balearica | Streptomyces viridobrunneus |
| Bacillus algicola | Glycomyces mayteni | Pseudomonas cichorii | Streptomyces werraensis |
| Bacillus alkalogaya | Halomonas hydrothermalis | Pseudomonas cinnamophila | Thiomonas intermedia |
| Bacillus carboniphilus | Halomonas johnsoniae | Pseudomonas fluorescens | Thiomonas perometabolis |
| Bacillus cohnii | Halomonas sinaiensis | Pseudomonas frederiksbergensis | Vagococcus salmoninarum |
| Bacillus djibelorensis | Halomonas variabilis | Pseudomonas mandelii | Varibaculum cambriense |
| Bacillus humi | Halothiobacillus halophilus | Pseudomonas metavorans | Vibrio furnissii |
| Bacillus infantis | Helcococcus kunzii | Pseudomonas moraviensis | Vibrio porteresiae |

Bacillus isabeliae
Bacillus koreensis
Bacillus krulwichiae
Bacillus methanolicus
Bacillus shackletonii
Bacillus siralis
Bacteroides fluxus
Bacteroides oleiciplenus
Bartonella weissi
Beijerinckia derxii
Bifidobacterium animalis
Bifidobacterium thermacidophilum
Bifidobacterium thermophilum
Bradyrhizobium elkanii
Brenneria quercina
Burkholderia mallei
Burkholderia sabiae
Candidatus Blochmannia
Candidatus Blochmannia herculeanus
Candidatus Blochmannia rufipes
Candidatus Contubernalis alkalaceticum
Candidatus Liberibacter africanus
Candidatus Phlomobacter fragariae
Candidatus Phytoplasma
Candidatus Phytoplasma pini
Candidatus Phytoplasma prunorum
Candidatus Protochlamydia
Candidatus Protochlamydia amoebophila

Hydrocarboniphaga effusa
Hymenobacter chitinivorans
Hyphomonas oceanitis
Jeotgalicoccus psychrophilus
Labrys wisconsinensis
Lactobacillus equi
Lactobacillus equicursoris
Lactobacillus fabifermentans
Lactobacillus oris
Lactobacillus pantheris
Lactobacillus plantarum
Lactococcus garvieae
Lampropedia hyalina
Leptothrix cholodnii
Leptothrix discophora
Leucobacter aridicollis
Leuconostoc pseudomesenteroides
Limnohabitans planktonicus
Loktanella koreensis
Lysinibacillus macroides
Lysinibacillus xylanilyticus
Lysobacter koreensis
Metallosphaera hakonensis
Methylobacterium fujisawaense
Methylobacterium persicinum
Methylophaga alcalica
Microbacterium laevaniformans
Moraxella catarrhalis

Pseudomonas mosselii
Pseudomonas oleovorans
Pseudomonas proteolytica
Pseudomonas putida
Pseudomonas rhodesiae
Pseudomonas savastanoi
Pseudomonas syringae
Pseudomonas tremae
Pseudomonas veronii
Pseudoxanthomonas sacheonensis
Psychrobacter adeliensis
Psychrobacter alimentarius
Psychrobacter celer
Psychrobacter cibarius
Psychrobacter cryohalolentis
Psychrobacter faecalis
Psychrobacter fozii
Psychrobacter glacincola
Psychrobacter immobilis
Psychrobacter marincola
Psychrobacter proteolyticus
Psychrobacter pulmonis
Psychrobacter submarinus
Ramlibacter tataouinensis
Rheinheimera chironomi
Rheinheimera soli
Rheinheimera texasensis
Rhizobium pisi

Virgibacillus kekensis
Virgibacillus sediminis
Weissella koreensis
Weissella soli
Xanthomonas sacchari
Yersinia mollaretii
Yersinia ruckeri
Zhihengliuella halotolerans

Candidatus Scalindua brodae
Candidatus Tammella caduceiae
Capnocytophaga leadbetteri
Capnocytophaga ochracea
Caulobacter crescentus
Caulobacter tundrae
Cetobacterium ceti
Chryseobacterium aquifrigidense
Chryseobacterium indologenes
Chryseobacterium ureilyticum
Clostridium neonatale
Clostridium papyrosolvens
Clostridium thermobutyricum
Corynebacterium casei
Corynebacterium glucuronolyticum
Corynebacterium pseudodiphtheriticum

Morganella morganii
Myroides injenensis
Myroides odoratimimus
Neisseria subflava
Nitratireductor kimnyeongensis
Nitrobacter hamburgensis
Nocardioides plantarum
Novosphingobium aromaticivorans
Novosphingobium capsulatum
Paenibacillus pini
Paracoccus carotinifaciens
Paracoccus marcusii
Paracoccus methylutens
Paracoccus niistensis
Paraprevotella xylaniphila

Rhodobaca bogoriensis
Rhodococcus baikonurensis
Rhodocyclus purpureus
Rhodoferax ferrireducens
Rhodovulum euryhalinum
Rothia aeria
Rummeliibacillus pycnus
Sanguibacter keddieii
Serratia symbiotica
Shewanella baltica
Shewanella frigidimarina
Shewanella glacialipiscicola
Shewanella livingstonensis
